# Single-pixel epiretinal stimulation with a wide-field and high-density retinal prosthesis for artificial vision

**DOI:** 10.1101/2020.08.21.261461

**Authors:** Naïg Aurelia Ludmilla Chenais, Marta Jole Ildelfonsa Airaghi Leccardi, Diego Ghezzi

## Abstract

Retinal prostheses hold the promise of restoring artificial vision in profoundly and totally blind people. However, a decade of clinical trials highlighted quantitative limitations hampering the possibility to reach this goal. A key obstacle to suitable retinal stimulation is the ability to independently activate retinal neurons over a large portion of the subject’s visual field. Reaching such a goal would significantly improve the perception accuracy in the users of retinal implants, along with their spatial cognition, attention, ambient mapping and interaction with the environment. Here we show a wide-field, high-density and high-resolution photovoltaic epiretinal prosthesis for artificial vision. The prosthesis embeds 10,498 physically and functionally independent photovoltaic pixels allowing for both wide retinal coverage and high-resolution stimulation. Single-pixel illumination reproducibly induced network-mediated responses from retinal ganglion cells at safe irradiance levels. Furthermore, the prosthesis enables a sub-receptive field response resolution for retinal ganglion cells having a dendritic tree larger than the pixel’s pitch. This approach could allow the restoration of mid-peripheric artificial vision in patients with retinitis pigmentosa.

---

Visual prostheses are artificial devices used to revert blindness^1–3^: a medical condition affecting more than 39 million people worldwide (World Health Organization). Over the years, several devices and solutions were proposed^4–9^, but so far retinal prostheses showed the best performances in patients affected by outer retinal layer dystrophies such as retinitis pigmentosa^10,11^ and age-related macular degeneration^12^. Indeed, the past ten years saw tremendous improvements from clinical^10,11^, pre-clinical^13–15^ and technological^16–18^ perspectives. Retinal implants mostly addressed blind patients with retinitis pigmentosa, a set of inherited retinal dystrophies with a prevalence of about 1:4,000 individuals, causing the progressive loss of retinal photoreceptors, the constriction of the visual field and eventually blindness^19^. Patients implanted with retinal prostheses were able to localise and identify letters or objects, and perform orientation tasks^20–22^. Despite the effort of the research community and the enthusiasm of the patients, the majority of the latter ceased using their implant in the first to the third year following their surgery^23^. Furthermore, one-third of the users of the Argus® II epiretinal prosthesis (the most implanted so far) declared that the device had a neutral impact on their quality of life after three years^24^. This discouragement can be attributed to quantitative limitations of retinal prostheses in daily use^23^.

Retinal prostheses approved by regulatory agencies provided at best a visual angle of 20 degrees (Argus® II^4^) and despite already being an incredible achievement in medical technology, it does not allow for safe and independent navigation in open spaces with obstacles and moving objects^25,26^. Independent mobility is of primary importance to increase the quality of life in profoundly blind patients with retinitis pigmentosa. Additionally, the coarse visual resolution offered by the device (i.e. for the Argus® II^4^, 6 × 10 electrodes with a 525-µm pitch), in combination with the small visual angle, provides little help in daily tasks involving object identification and recognition. Last, patients reported that the use of retinal prostheses is cognitively exhausting due to the constant need of space decomposition^23^: because of the limited visual angle, the users are instructed to move their head and body to scan the environment. Scanning implies a constant visual decomposition and mental reconstruction of the visual scene, often guided by a complex pairing of the coarse visual information with audio-tactile cues^23^. Studies under simulated prosthetic vision identified a visual angle of 30 degrees as the minimal requirement to efficiently complete everyday mobility and manipulation tasks^27–30^. However, this number might underestimate the real needs of retinal prostheses’ users, which exhibit poor performance in those tasks, due to the perceptual learning and behavioural adaptation issues to the new form of partial and spatially fractioned artificial vision^31,32^. The visual angle is a significant bottleneck preventing patients from achieving an efficient performance and a sustainable experience. New retinal prostheses should overcome these challenges and restore a large enough visual angle fitting the natural scanning via eye movements to provide a helpful and valuable visual aid to patients with retinitis pigmentosa. Wide-field retinal prostheses enabling theoretical visual angles larger than 30 degrees were recently proposed and tested in preclinical studies to meet these requirements^16,33^.

Nevertheless, the visual angle is not the only barrier for retinal prostheses’ users. Object identification and recognition require devices able to provide enough resolution. Wide-field arrays were so far designed for epiretinal placement only since very large subretinal implants might encounter considerable difficulty in the surgical placement and represent a high risk of retinal detachment^34–36^. However, clinical trials showed that the best visual resolution was achieved using subretinal prostheses. The highest visual acuities reported to date, as measured with Landolt-C test, were 20/460 (logMAR 1.37) with the subretinal implant PRIMA^12^ and 20/546 (logMAR 1.43) with the subretinal implant Alpha-AMS^37^. Grating acuities reported in the literature range from 20/1260 with the epiretinal Argus® II implant^38^ to 20/364 with the subretinal Alpha-AMS implant^10^. The inadequate performance of epiretinal prostheses like the Argus® II can be attributed to two factors: on the one hand, the implantable pulse generator, the transscleral cable and the feedlines in the array strongly limits the number and density of the electrodes. On the other hand, the activation of the nerve fibre layer reduces the spatial resolution of epiretinal stimulation.

In order to further address the quantitative limitations aforementioned, we designed and characterised a wide-field curved organic photovoltaic epiretinal prosthesis with a very high pixel density. This high-density implant was conceived to offer a relevant resolution for the patient’s needs through epiretinal network-mediated stimulation, thus overcoming the direct activation of the nerve fibre layer, and first and foremost a large visual angle with minimal need of head scanning. However, the high pixel density of the prosthesis does not necessarily correlate with a high response resolution at the retinal ganglion cell (RGC) level^39–41^, due to the spatial interconnectivity of the retinal network. Therefore, we investigated ex vivo the response resolution provided by this high-density retinal prosthesis. Our results showed that the wide-field and high-density organic photovoltaic epiretinal prosthesis allows for single-pixel stimulation of RGCs at safe irradiance levels. Also, sub-portions of the RGC receptive field (RF) are recruited through the photovoltaic pixels, which confirms the high-resolution of the network-mediated epiretinal stimulation. These results demonstrated that an epiretinal prosthesis could achieve a high spatial resolution in retinal stimulation, together with a wide visual angle, which together would be a substantial step forward for artificial vision.

## Results

### High-density retinal prosthesis

POLYRETINA is a wide-field high-density epiretinal prosthesis which contains 10,498 photovoltaic pixels (80-µm diameter, 120-µm pitch) distributed with a density of 79.1 pixels mm^−2^ over an active area of 13 mm in diameter (**Fig. 1a,b**). Once bonded to its curved flexible support, the active area is slightly stretched to 13.4 mm, and the prosthesis covers a visual angle of about 43 degrees (750 mrad). Compared to the previous POLYRETINA design^16^, the number of pixels and their density was increases, together with two other technical improvements. First, titanium (Ti) electrodes were coated with a layer of titanium nitride (TiN) to enhance the stimulation efficiency while keeping a safe capacitive stimulation. Second, the polymer-based layers below each cathode were patterned to generate physically independent photovoltaic pixels (**Fig. 1c**) and to avoid having cracks between rigid platforms made out of SU-8 (**Fig. 1b**).

**Figure 1.**
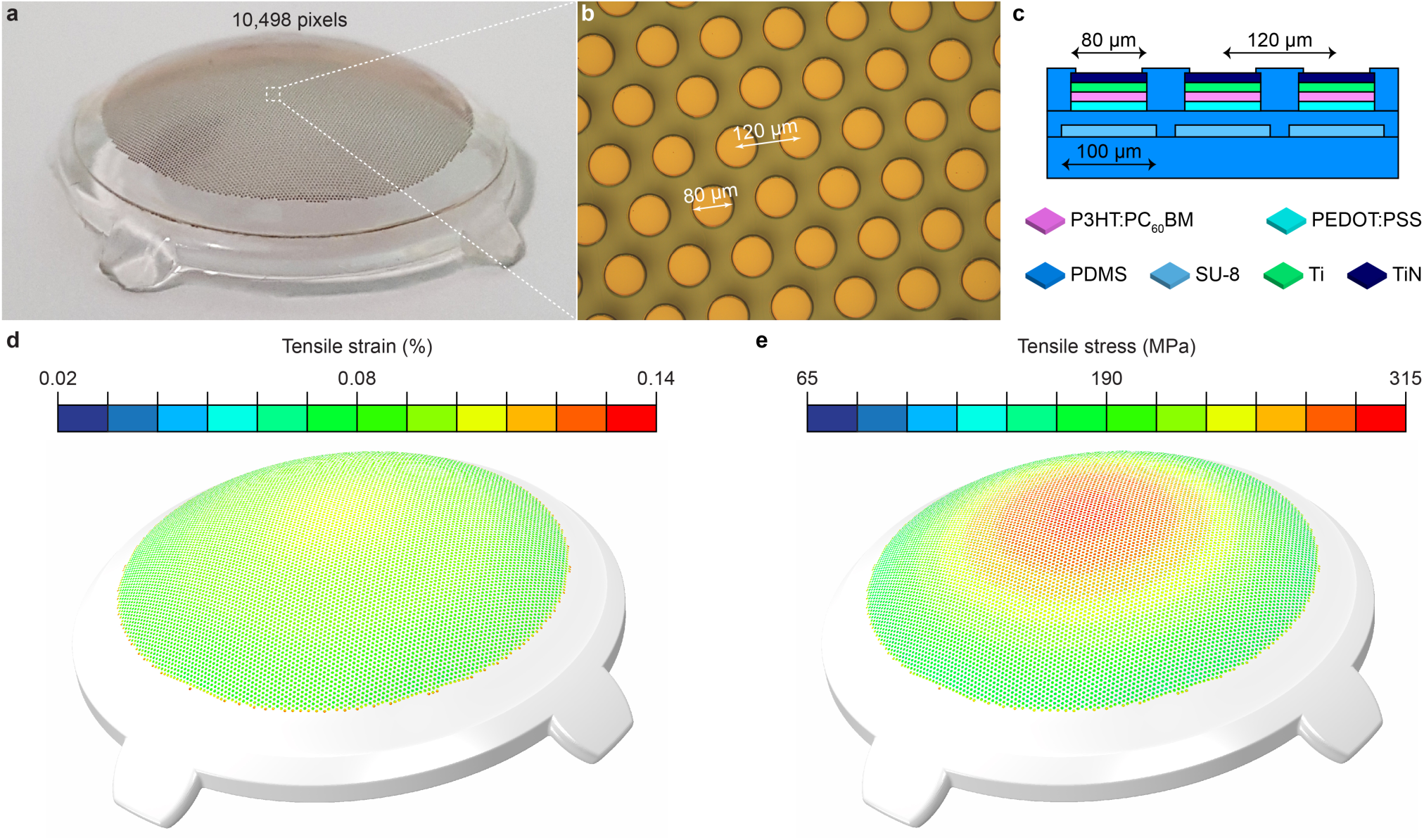
High-density POLYRETINA device. **a**, Picture of a fabricated high-density POLYRETINA prosthesis with 10,498 photovoltaic pixels. **b**, Magnified view of the photovoltaic pixels having a diameter of 80 µm and a pitch of 120 µm. **c**, Sketch of the cross-section structure of the POLYRETINA photovoltaic interface before bonding to the hemispherical dome. The layer thicknesses are as follow: base PDMS layer: 50 μm; SU-8 platforms: 6 μm; second PDMS layer embedding SU-8 platforms: 15 µm; PEDOT:PSS: 50 nm, P3HT:PCBM: 100 nm, Ti-TiN: 80-60 nm, final PDMS layer: 4 μm. PDMS: polydimethylsiloxane; PEDOT: poly(3,4-ethylenedioxythiophene); PSS: poly(styrenesulfonate); P3HT: regioregular poly(3-hexylthiophene-2,5-diyl); PC_60_BM: [6,6]-phenyl-C61-butyric acid methyl ester; Ti: titanium; TiN: titanium nitride. **d**, Tensile strain simulated at the level of TiN. **e**, Tensile stress simulated at the level of TiN.

The fabrication of a high-density array brings on several challenges. First, the higher the pixel density, the higher the risk that the pixels would crack during the hemispherical shaping of the device. We performed finite element analysis simulations to estimate the level of tensile stress and strain occurring onto the cathodes during hemispherical shaping (**Fig 1d,e**). The TiN coating reduced the tensile strain from −0.55% (Ti pixels) to −0.13% (TiN-coated pixels) and the tensile stress from 574.8 MPa (Ti) to 310.9 MPa (TiN). The reduction of tensile stress during hemispherical shaping further protects the metal cathodes (**Fig. 1b**).

Second, a higher pixel density might induce crosstalk during stimulation with neighbouring pixels. To rule out this possibility, we measured the radial voltage spreading (**Fig. 2a**) in three directions (D1, D2 and D3 in **Fig. 2b**) using a glass microelectrode upon pulsed illumination of a single-pixel (560 nm, 10 ms). The minimum irradiance level required to activate RGCs ex vivo is about one hundred of µW mm^−2^ for large-field illumination^16^; yet, to exclude crosstalk even at very high irradiance levels, we performed the experiment at 22.65 mW mm^−2^, the maximal irradiance attainable by the illumination system. For each direction, the voltage generated from the pixel was measured in several points at increasing distance from the illuminated pixel (red points in **Fig. 2b**) and interpolated in a two-dimensional colour map (**Fig. 2c**). The voltage generated by the illumination of a single-pixel remained localised above the pixel. In order to ensure that neighbouring pixels do not induce crosstalk, we repeated the experiment activating one pixel (**Fig. 2d**, left), one pixel with one surrounding corona of pixels (seven pixels; **Fig. 2d**, middle left), one pixel with two surrounding coronas of pixels (nineteen pixels; **Fig. 2d**, middle right), or the two surrounding coronas of pixels with the central pixel off (eighteen pixels; **Fig. 2d**, right). For each condition, the normalised voltage profiles in the three principal directions were averaged. The average plot of the voltage profile showed that the voltage generated by each pixel is sharply discriminated from the one of neighbouring pixels and does not show a voltage summation effect, in all the configurations tested (**Fig. 2e**). Even in the extreme case where the central pixel is off while the surrounding eighteen pixels are on, there is very high contrast in the voltage drop between the central pixel and the neighbouring ones (**Fig. 2e**, right), although a small residual potential is present also onto the central pixel. These results show that the pixels are electrically independent (i.e. no crosstalk). However, it must be noted that the voltage measures were taken close to the device’s surface (2-5 µm distance). Such a sharp discrimination of the voltage profile might be reduced at larger distances from the array, where RGCs and bipolar cells (BCs) are located.

**Figure 2.**
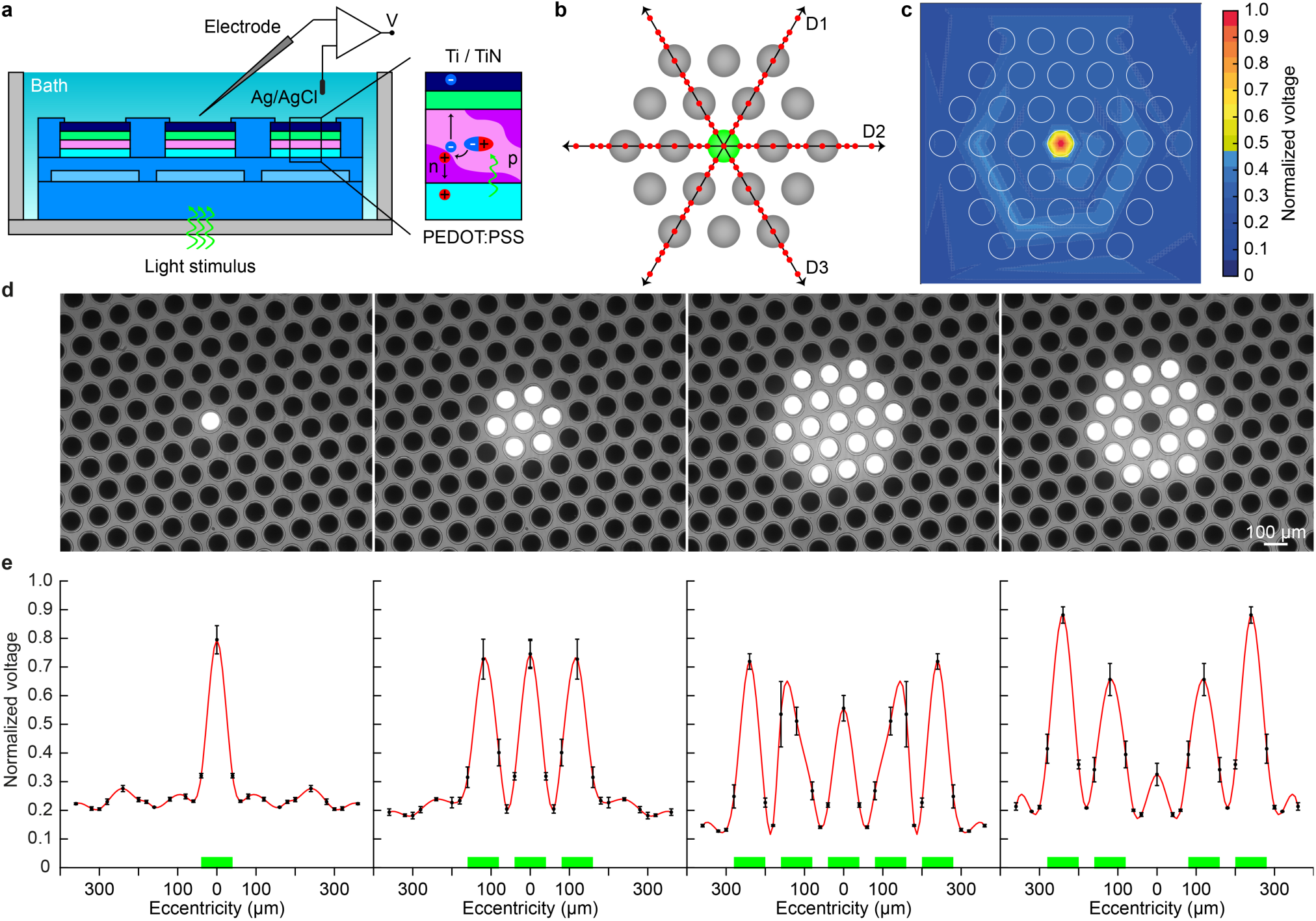
Stimulation selectivity of the photovoltaic pixel. **a**, Sketch of the experimental set-up to measure the voltage spreading. The insert shows the photovoltaic transduction mechanism. A photon is absorbed by the P3HT (p-type semiconductor, electron donor) and an exciton is formed. The exciton travels until it reaches the interface between P3HT and PCBM (n-type semiconductor, electron acceptor) and dissociates. The electron is attracted towards the cathode (Ti/TiN), and the hole is attracted towards the anode (PEDOT:PSS) because of their work function levels. **b**, Sketch of the experimental methodology. The green dot corresponds to the illuminated pixel (560 nm, 10 ms, 22 mW mm^−2^). The grey ones represent the surrounding pixels. The voltage was measured in 25 positions (red dots) for each direction (D1, D2, and D3). **c**, Voltage spreading colour map generated by interpolating the experimental measures with a triangulation-based linear interpolation. For each data point, 10 consecutive recordings were averaged and the voltage peaks were normalised to the maximal value obtained in the whole experiment. The white circles show the location of the pixels. **d**, Pictures of the four stimulation patterns: central on (left), central on and one corona on (middle left), central on and two coronas on (middle right), and central off and two coronas on (right). The light spots are visible (brighter area). **e**, Normalised voltage profiles obtained for the four illumination patterns (mean ± s.e.m.; *n* = 4 prostheses). For each prosthesis, the normalised data from the 3 directions were averaged. The red line shows a Gaussian fitting, and the green bars beneath corresponds to the active pixels.

Third, high-density retinal prostheses would represent a useful advancement only if stimulation of RGCs can be achieved by single-pixel illumination. Thus, TiN was coated on top of the pixels to increase their stimulation efficiency. Using Kelvin Probe Force Microscopy (KPFM), we evaluated the changes in the surface potential generated at the cathode upon illumination (560 nm, 60 s, 0.9 mW mm^−2^) with and without the TiN coating (**Fig. 3a**). The irradiance level was set to 0.9 mW mm^−2^ since our previous results showed a saturation of the RGC response beyond this value^16^. TiN-coated pixels showed a statistically significant higher change in the surface potential compared to Ti pixels (**Fig. 3b**; p = 0.0083, two-tailed unpaired t-test). Next, we measured the photo-current (PC) and the photo-voltage (PV) generated by the pixels upon illumination (565 nm, 10 ms) at increasing irradiance levels. We fabricated chips embedding 6 pixels, each of them connected to a contact pad to measure the signal at the cathode against a platinum reference electrode immersed in saline solution (**Fig. 3c**). The mean PC density (PCD) and PV were both higher for TiN-coated pixels upon illumination at increasing irradiance levels (**Fig. 3f,g**). We further evaluated the PCD and the PV at the representative irradiance level of 0.9 mW mm^−2^ (**Fig. 3h**): a statistically significant difference between Ti and TiN-coated pixels was found for both the PCD (p = 0.0288, two-tailed unpaired t-test) and the PV (p < 0.0001, two-tailed unpaired t-test). The surface area of the cathodes was measured with an atomic force microscope (AFM) over an area of 500 × 500 nm^2^ (**Fig. 3i,j**). On average (*n* = 3 pixels), TiN-coated pixels showed a statistically higher surface area compared to Ti pixels (p = 0.0024, two-tailed unpaired t-test).

**Figure 3.**
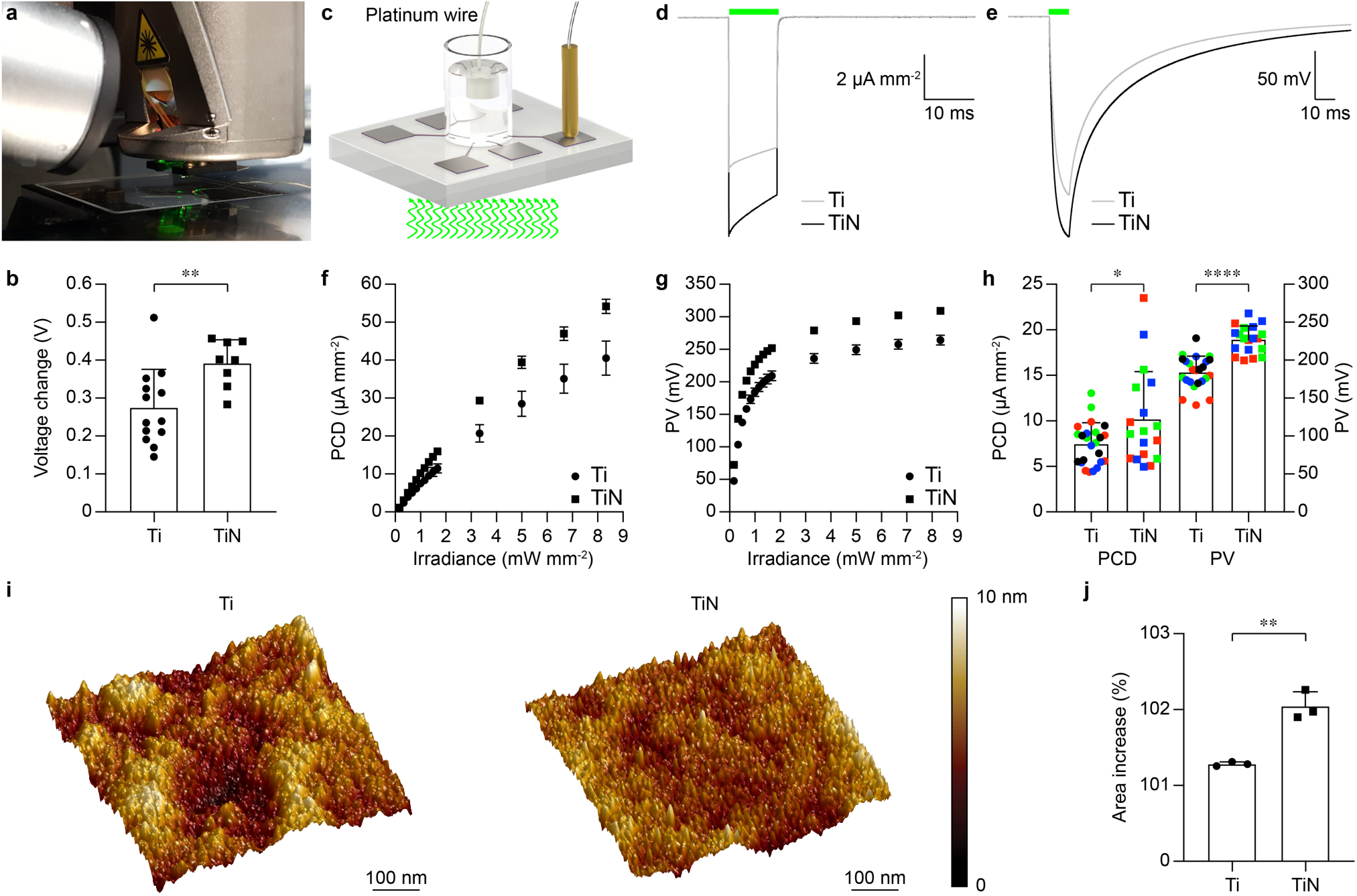
Optoelectronic characterisation of the photovoltaic pixel. **a**, Picture of the KPFM set-up. **b**, Surface voltage changes obtained with Ti (circles) and TiN-coated (squares) cathodes. Each bar is the mean (± s.d.) of the measures from *n* = 13 Ti pixels and *n* = 8 TiN-coated pixels. **c**, Drawing of the experimental set-up for the measure of the PC and PV; the light pulse comes from the bottom. **d**,**e**, Mean PCD (**d**) and PV (**e**) measures obtained from Ti (grey) and TiN-coated (black) pixels upon illumination (565 nm, 10 ms, 0.9 mW mm^−2^). For Ti, *n* = 24 pixels from 4 chips were averaged; for TiN, *n* = 18 pixels from 3 chips were averaged. **f**,**g**, Mean (± s.e.m) PCD (**f**) and PV (**g**) amplitudes quantified at increasing irradiance levels (565 nm, 10 ms) for Ti pixels (circles; *n* = 24 pixels from 4 chips) and TiN-coated pixels (squares; *n* = 18 pixels from 3 chips). **h**, Mean (± s.e.m) PCD and PV amplitudes quantified at 0.9 mW mm^−2^ for Ti (*n* = 24 pixels from 4 chips) and TiN-coated (*n* = 18 pixels from 3 chips) pixels. **i**, AFM images of the Ti and Ti/TiN surfaces. The colour bar shows the surface roughness. **j**, Mean (± s.d.) percental increase of the surface area compared to the nominal flat area (500 × 500 nm^2^).

These results confirmed that the photovoltaic pixels are physically and functionally independent. The coating with TiN reduced the mechanical stress of the pixels and increased their photovoltaic performance by likely reducing the electrode-electrolyte impedance, increasing the interface capacitance and reducing the parasitic resistances of the photovoltaic pixel. These results open up the possibility of high-resolution single-pixel stimulation of RGCs.

### Single-pixel stimulation efficiency of titanium nitride photovoltaic pixels

We subsequently evaluated whether the increased photovoltaic performances of TiN-coated pixels translated into a higher stimulation efficiency of RGCs. For the study, we used explanted retinas from the retinal degeneration 10 (rd10) mouse model, which is an established model for retinitis pigmentosa^42–44^. In agreement with our previous study^45^ and studies performed by other laboratories^46,47^, rd10 retinas beyond post-natal day (P) 60 can be considered light-insensitive. In order to ensure a proper exclusion of intrinsic light responses due to spared photoreceptors, the experiments in this work were performed in rd10 retinas at a very late stage of degeneration (mean age ± s.d.: 126.7 ± 15.1; **Tab. 3**). Also, both male and female mice were used to exclude any effect related to sex-related differences in the degeneration onset and progression in rd10 retinas (**Tab. 3**). Explanted retinas were layered in epiretinal configuration, and the prosthetic-evoked activity of RGCs was recorded via single-electrode extracellular recordings (**Fig. 4a**). Light pulses (560 nm, 10 ms) were delivered in a broad range of irradiance levels (0.9, 2.34, 6.24, 12.37, 17.68 and 22.65 mW mm^−2^) and the network-mediated medium-latency (ML) responses of RGCs to large-field (covering approximately 70 pixels) illumination (**Fig. 4b**) and single-pixel illumination (**Fig. 4c**) were compared. The quantification of the ML spiking activity upon large-field illumination revealed that TiN-coated pixels elicited on average higher ML spiking activity than Ti pixels (**Fig. 4d**). Moreover, in both conditions, the first irradiance tested (0.9 mW mm^−2^) elicited a statistically significant ML spiking activity higher than the basal activity computed without light (Ti: p = 0.0088; TiN: p < 0.0001; two-tailed unpaired t-test). When the illumination was switched to single-pixel (**Fig. 4e**), both Ti and TiN-coated pixels also evoked a statistically significant ML spiking activity at the first irradiance tested (0.9 mW mm^−2^; Ti: p = 0.0378; TiN: p = 0.0062; two-tailed unpaired t-test). This result revealed that both Ti and TiN-coated photovoltaic pixels induced RGC activity upon single-pixel illumination.

**Figure 4.**
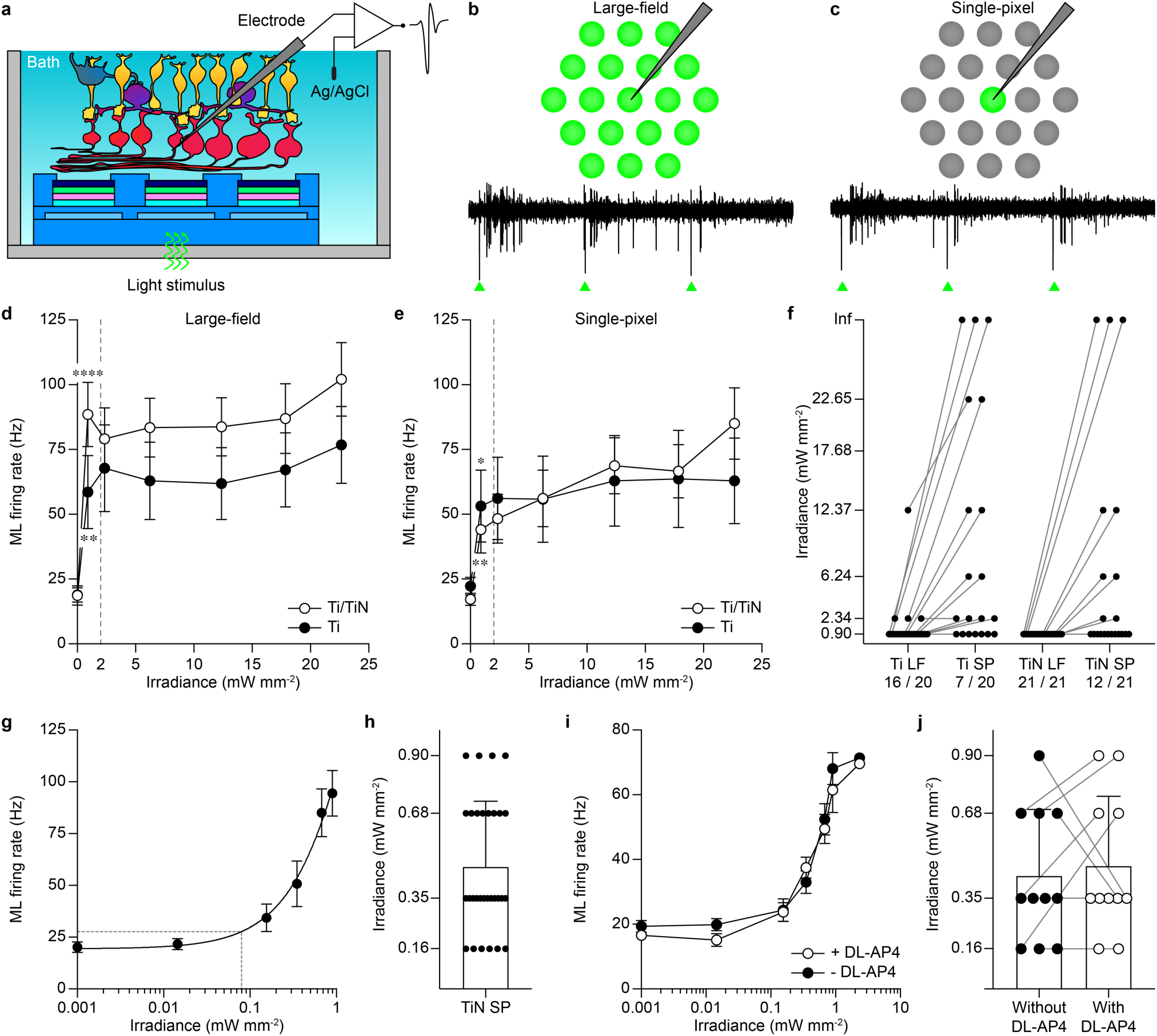
Single-pixel stimulation of retinal ganglion cells. **a**, Sketch of the recording set-up. **b**,**c**, Representative responses from a retinal ganglion cell upon 3 consecutive light pulses (560 nm, 10 ms, 0.9 mW mm^−2^) with large-field illumination (**b**) and single-pixel illumination (**c**). **d**,**e**, Comparison of the stimulation efficacy during large-field (**d**) and single-pixel (**e**) illumination under increasing irradiances with Ti-based (*n* = 20 RGCs, mean ± s.e.m.) and TiN-based pixels (*n* = 21 RGCs, mean ± s.e.m.). **f**, Change in the activation threshold from large-field to single-pixel illumination for both Ti pixels (*n* = 20 RGCs) and TiN-coated pixels (*n* = 21 RGCs). The numbers for each column are the fraction of RGCs activated by a 10-ms light pulse of 0.9 mW mm^−2^. Inf means that the RGC does not show ML activity at any of the irradiance level tested. LF: large-field, SP: single-pixel. **g**, ML firing rate (*n* = 30 RGCs, mean ± s.e.m.) at low irradiance levels for TiN-coated pixels and single-pixel illumination. The black line is the second-order polynomial interpolation (R squared = 0.29). The grey dashed lines show the activation threshold computed as the irradiance eliciting 10% of the maximal ML firing rate at 0.9 mW mm^−2^. **h**, Activation threshold for each RGCs with single-pixel illumination and TiN-coated pixels (mean ± s.d.). **i**, ML firing rate (*n* = 11 RGCs, mean ± s.e.m.) at low irradiance levels for TiN-coated pixels and single-pixel illumination before and after the application of DL-AP4. **j**, Activation threshold for each individual RGC with single-pixel illumination and TiN-coated pixels before and after the application of DL-AP4 (mean ± s.d.).

**Table 3.** Animal groups. Sex and age of the rd10 mice used in the study.

| Ti vs TiN comparison |  |  | TiN irradiance threshold |  | TiN RF |  | TiN pixel switch |  |
| --- | --- | --- | --- | --- | --- | --- | --- | --- |
| Fig. 4b-f |  |  | Fig. 4g,h |  | Fig. 5 |  | Fig. 6 |  |
| Sex | Age | Material | Sex | Age | Sex | Age | Sex | Age |
| male | 133 | Ti | male | 115 | female | 137 | male | 108 |
| male | 141 | Ti | male | 123 | male | 117 | male | 130 |
| male | 108 | Ti | female | 127 | male | 119 | male | 108 |
| female | 116 | Ti | female | 134 | male | 148 | male | 108 |
| female | 114 | Ti | Pharmacology<br>Fig. 4g,h,i,j |  | male | 123 |  |  |
| male | 117 | Ti |  |  | female | 127 |  |  |
| male | 142 | Ti | male | 115 | female | 134 |  |  |
| male | 146 | Ti | male | 107 | male | 135 |  |  |
| male | 148 | Ti | male | 108 | male | 138 |  |  |
| female | 142 | TiN | female | 147 | male | 139 |  |  |
| male | 108 | TiN | female | 143 |  |  |  |  |
| female | 96 | TiN |  |  |  |  |  |  |
| male | 119 | TiN |  |  |  |  |  |  |
| male | 146 | TiN |  |  |  |  |  |  |
| male | 148 | TiN |  |  |  |  |  |  |

Next, we quantified the fraction of RGCs that could be activated with 10-ms pulses at 0.9 mW mm^−2^ under both large-field and single-pixel stimulation. For Ti pixels and large-field stimulation, 16 out of 20 RGCs showed ML responses at 0.9 mW mm^−2^, or in other words, exhibited an activation threshold lower or equal to 0.9 mW mm^−2^. 3 out of 20 RGCs showed activation at 2.34 mW mm^−2^, and 1 out of 20 RGCs showed activation at 12.37 mW mm^−2^. Switching to single-pixel illumination, only one-third of the RGCs (7 out of 20) preserved ML activation upon illumination at 0.9 mW mm^−2^. For TiN-coated pixels, all RGCs (21 out of 21) show ML activity upon large-field illumination at 0.9 mW mm^−2^, and still more than half RGCs (12 out of 21) when the illumination was switched to single-pixel. This result shows the higher efficiency in single-pixel retinal stimulation of TiN-coated photovoltaic pixels compared to Ti pixels. Noteworthy, this increase in efficiency cannot be attributed to differences in the animals used. For Ti pixels, nine animals were used (129.4 ± 15.6, mean age ± s.d), of which 7 males (77.8%) and 2 females (22.2%). For TiN-coated pixels, six animals were used (126.5 ± 22.0, mean age ± s.d), of which 4 males (66.7%) and 2 females (33.3%). Also, the mice’s age was not statistically different among the two groups (p = 0.77, two-tailed unpaired t-test).

In a second set of cells (*n* = 30 RGCs) exhibiting ML response upon single-pixel illumination of TiN-coated pixels at 0.9 mW mm^−2^, we determined the threshold for activation using lower irradiance levels (0.014, 0.16, 0.35, 0.68, and 0.9 mW mm^−2^). The average ML firing rates upon single-pixel illumination increased as a function of the irradiance (**Fig. 4g**) with an activation threshold of about 79 µW mm^−2^, obtained as the irradiance level providing 10% of the maximal ML firing rate measured at 0.9 mW mm^−2^. This result shows that the responsivity of the RGCs can be modulated as a function of the irradiance level. However, we observed that the single-pixel activation threshold for individual RGCs is variable among the irradiance levels tested (**Fig. 4h**), and more than half (18 out of 30) of the RGCs exhibited a ML response threshold lower or equal to 0.35 mW mm^−2^. While the population threshold was estimated to be 79 µW mm^−2^, only 6 out of 30 cells showed ML responses at 160 µW mm^−2^. The disparity of the network-mediated ML activation thresholds can be related to the location of the cell’s RF compared to the position of the illuminated pixel. Due to the recording method, the cell’s soma or the upstream BCs cannot be precisely located. Therefore, RGCs having their RF centred over a pixel might have a lower threshold than those RGCs eccentric to the pixel because of the very limited lateral spreading of the photovoltaic stimulus. Last, we evaluated in a subset of RGCs (*n* = 11, TiN-coated pixels) the ML responsivity with and without the application of a broad spectrum glutamatergic synaptic antagonist (DL-AP4, 250 μM l^−1^ ; No. 0101, Tocris Bioscience), which blocks the synaptic input of ON BCs^48^ (**Fig. 4i**). The ML response curve was not altered by the introduction of the antagonist, thus excluding any contribution from potential spared photoreceptors to the ML responses recorded upon photovoltaic stimulation. Also, the single-pixel activation threshold measured in individual RGCs was not statistically different after DL-AP4 application (**Fig. 4j**; p = 0.68, two-tailed paired t-test).

These results confirmed that the network-mediated stimulation of RGCs in epiretinal configuration is achieved with single-pixel illumination at irradiance levels largely below the maximum permissible exposure (MPE) limit for retinal safety, which for POLYRETINA varies between 8.32 and 2.08 mW mm^−2^ respectively for 5 and 20 Hz illumination rate.

### Photovoltaic receptive fields

Using TiN-coated pixels and single-pixel illumination at 0.9 mW mm^−2^, we quantified the number of pixels able to activate a RGC (**Fig. 5**). The 19 neighbouring pixels around the recording electrode were successively illuminated (560 nm, 10 ms, 0.9 mW mm^−2^) according to a counter-clockwise pattern (**Fig. 5a**). The network-mediated ML activities elicited by the illumination of each pixel were mapped to the pixel coordinates, and the photovoltaic RFs of the RGCs were fitted with a two-dimensional gaussian model. Each RF diameter was then calculated as the average between the horizontal and vertical standard deviation of its two-dimensional activation map. The majority of the recorded RGCs (24 out of 31 cells) exhibited small RFs with a radius ranging from 34.5 to 142.5 µm (**Fig. 5d**). 5 out of 31 RGCs exhibited large RFs whose radius varied between 184.3 and 282.7 µm (**Fig. 5e**). 2 out of 31 RGCs exhibited elongated RFs, showing high responses to several aligned pixels (**Fig. 5f**). The activation maps of the RGCs could be clustered (Gaussian mixture model) into two populations (**Fig. 5b**), namely those exhibiting small or large RFs (RGCs with elongated RFs were excluded from the analysis). For clustering, the RFs were rotated so that the horizontal direction (x-axis) corresponds to the axis of maximal dispersion and the vertical direction (y-axis) corresponds to the dispersion in the orthogonal direction. The average RF diameter for each population was respectively 153.7 ± 26.1 µm and 335.5 ± 49.3 µm (mean ± s.e.m). In both the healthy and the rd10 mouse retinas, most of the RGCs exhibit functional RFs of a similar size (with a median cluster diameter between 119 and 280 µm)^49–51^ to the RGCs exhibiting a small photovoltaic RF. Also, populations of large α-RGCs with larger dendritic trees (up to 395 µm diameter) were reported in the mouse retina^50,52^, with a RF similar to the RGCs exhibiting a large photovoltaic RF. A statistical analysis revealed that RGCs with small RFs are indirectly stimulated through an average of three photovoltaic pixels (**Fig. 5c**). A pixel is considered to induce statistically significant activation of the RGC if the mean ML response, evaluated over ten repetitions, is statistically significantly higher than the cell background activity (two-tailed unpaired t-test, p < 0.05), which was calculated as the activity in the 100-ms pre-stimulus period averaged across all the illuminated pixels. Therefore, in mice retinas, the minimum response resolution offered by the high-density POLYRETINA for RGC activation is limited by the natural RF of the targeted cell (due to the network-mediated mechanism), rather than by the pixel density. Cells with very small RFs can be activated by one pixel only, and this number increases with the photovoltaic RF size. However, the resolution limit might also be affected by the actual centring of the cell and its RF compared to the photovoltaic pixels: a single RGC can be activated by multiple neighbouring pixels when its RF is centred in between the pixels.

**Figure 5.**
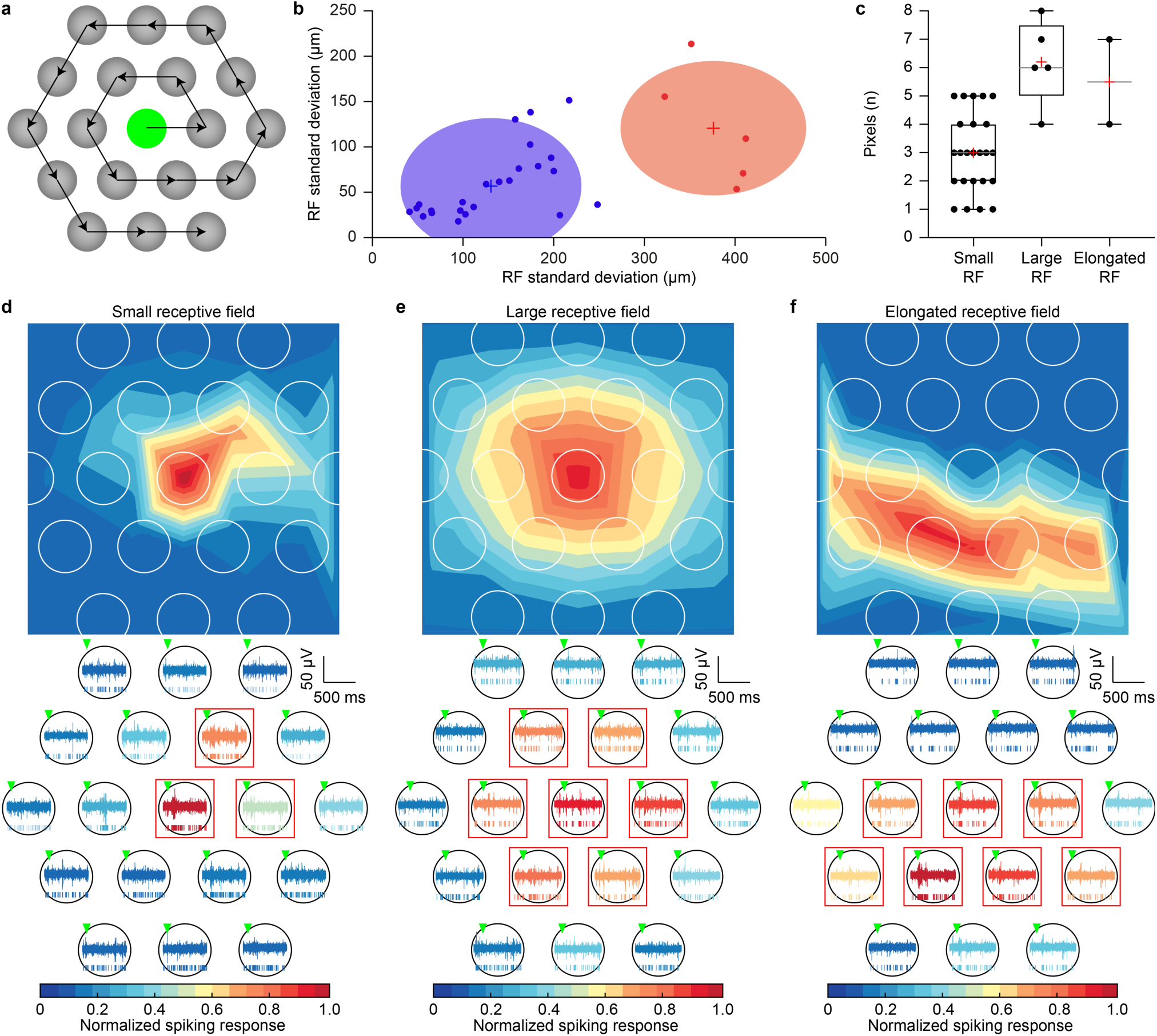
Photovoltaic epiretinal receptive fields. **a**, Sketch of the temporal pattern used for stimulation. Each of the 19 pixels centred around the recording location was successively illuminated following a counter-clockwise pattern (560 nm, 10 ms, 0.9 mW mm^−2^). The total illumination sequence was repeated for 10 consecutive sweeps. The receptive fields or activation maps were obtained by two-dimensional Gaussian approximation of the network-mediated ML responses elicited by single-pixel, averaged over sweeps, and normalized to the maximal responding pixel. **b**, Gaussian mixture model of the RF sizes over the small (*n* = 24 RGCs) and large RGC populations (*n* = 5 RGCs). The average RF diameter of the small and large cells types are respectively of 153.7 ± 26.1 µm and 335.5 ± 49.3 µm (mean ± s.e.m.). **c**, Quantification of the number of pixels able to induce statistically significant (p < 0.05) ML activation in the recorded RGC. The horizontal grey line is the median, the red plus is the average and the the boxes extend from the 25^th^ to 75^th^ percentiles. **d**,**e**,**f**, Photovoltaic RFs from three individual RGCs classified as small RF cell (**d**), large RF cell (**e**), and elongated RF cell (**f**). The bottom panels show raw electrophysiological recordings and raster plots from the same cells for each SP illumination (560 nm, 10 ms, 0.9 mW mm^−2^, first sweep). The red boxes show the pixels inducing a statistically significant activation of the recorded RGC.

### Spatial resolution of single-pixel stimulation

According to previous studies^53,54^, the retina desensitises upon repetitive network-mediated stimulation, and the RGC spiking response decays proportionally to the stimulation frequency. Upon repetitive stimulation from the same pixel at 5 Hz (560 nm, 10 ms, 0.9 mW mm^− 2^, 10 pulses) the desensitisation can be observed already at the second light pulse and it reaches a steady-state after the fourth pulse (**Fig. 6a-c**). Taking advantage of this ineluctable desensitisation process, we investigated the effect of a simple pattern reversal on the RGC response to assess whether adjacent pixels are able to recruit a single RGC selectively. We repeatedly stimulated RGCs in an alternating pattern with the two most responding pixels for each cell within their RFs (**Fig. 6d,e**). Upon repetitive stimulation from the same pixel at 5 Hz (560 nm, 10 ms, 0.9 mW mm^−2^, 5 pulses) the desensitisation can be observed already at the second light pulse (**Fig. 6f**) with a 27.3 ± 11.2% drop in the ML firing rate (mean ± s.e.m.) compared to the first pulse response (p = 0.0105, two-tailed paired t-test) and a τ = 0.93 ± 0.14 s decay constant (mean ± s.e.m.) over the five consecutive pulses. However, the ML response is recovered at the pixel switch as strongly as the response to the first pulse (p = 0.9054, two-tailed paired t-test). This result suggests that neighbouring pixels can independently target the same RGC through different portions of its RF, making it sensitive to sub-RF resolution pattern reversals.

**Figure 6.**
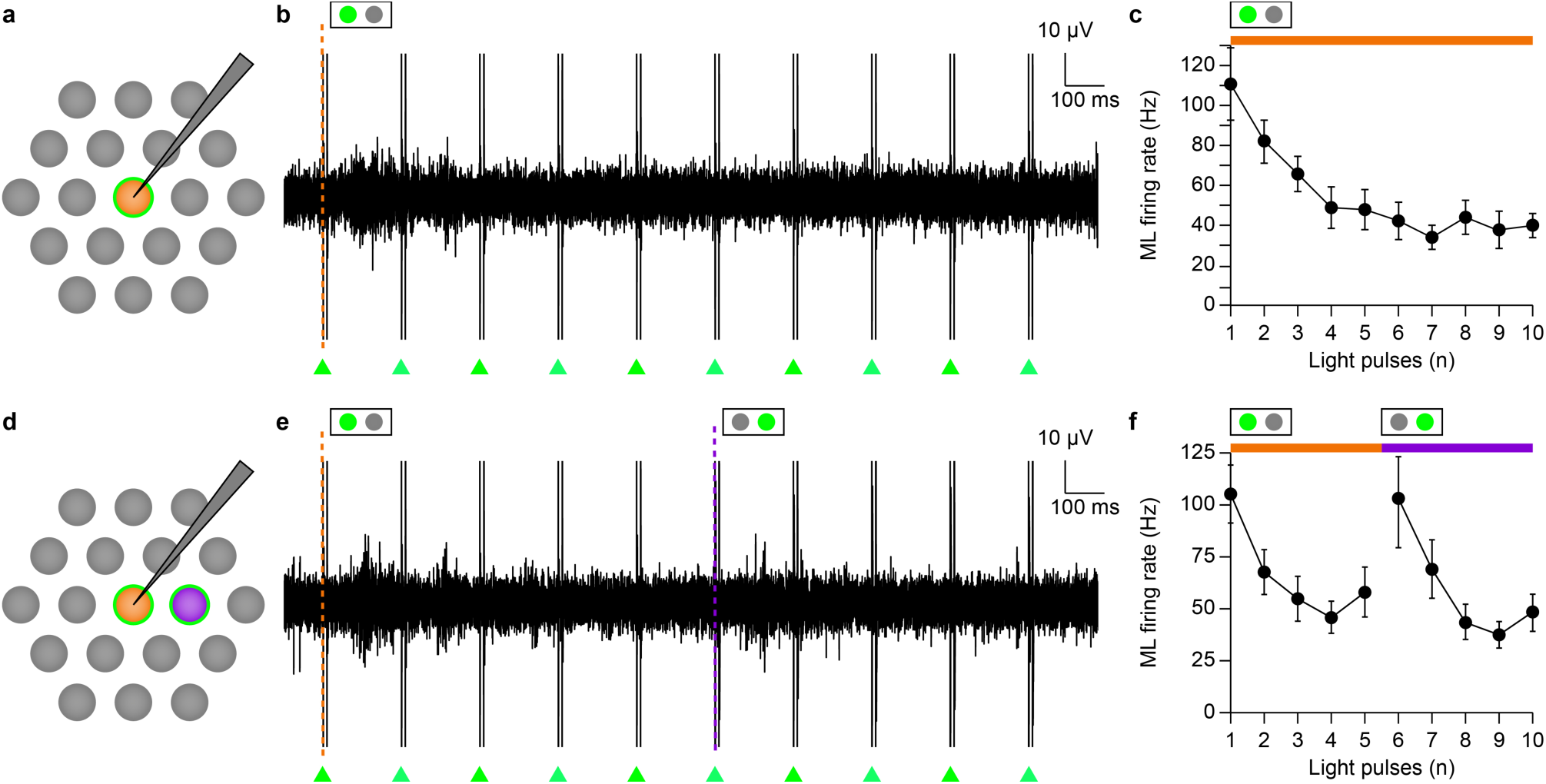
Spatial activation of the RGC layer under single-pixel switch stimulation. **a**, Sketch of the pattern used for stimulation without pixel switch. One pixel was successively illuminated at 5 Hz (560 nm, 10 ms, 0.9 mW mm^−2^) for 1 s. **b**, Raw electrophysiological recordings of a single RGC under 5 Hz continuous stimulation through one pixel **c**, ML firing rate (mean ± s.e.m.) over continuous stimulation through one pixel within a single RGC RFs (*n* = 12 RGCs). **d**, Sketch of the pattern used for stimulation with pixel switch. Each of the two-coloured pixels was successively illuminated at 5 Hz (560 nm, 10 ms, 0.9 mW mm^−2^) for 1 s, and the switched to the adjacent pixel at 1 Hz. **e**, Raw electrophysiological recordings of a single RGC under 5Hz continuous switch stimulation through two adjacent pixels **f**, ML firing rate (mean ± s.e.m.) over continuous switch stimulation through adjacent pixels within a single RGC RFs (*n* = 12 RGCs).

## Discussion

So far, the maximum number of electrodes embedded in retinal prostheses, as well as their overall density were limited by the use of implantable pulse generators, trans-scleral connections and feedlines in the array^36^. The introduction of the photovoltaic technique in the field of retinal prostheses has overcome all the problems mentioned above in a single step^55^. However, despite of this advancement, the small size, high stiffness and low conformability of many devices keep limiting the overall retinal coverage, and so the restored visual angle^14,15,56^. Conjugated polymers combined with stretchable substrates overcomes this last issue^16^; indeed, the POLYRETINA prosthesis enables at the same time photovoltaic retinal stimulation and wide coverage of the retinal surface. The results presented in this article also show that the high-density POLYRETINA allows for an epiretinal stimulation at high spatial resolution. The microscale patterning of conjugated polymers is an essential element to fabricate such high-density wide-field organic photovoltaic prosthesis. 10,498 physically and electrically independent pixels were manufactured into the array with a 120-µm pitch. The mechanical integrity of the device was preserved thanks to both the patterning of the conjugated polymers and the TiN coating, which reduces the tensile stress on the pixels. The mechanical compliance of the device allows the bonding of the high-density array over a large soft hemispherical dome in order to maintain close contact between the pixels and the retinal tissue over the central and peripheric retina. Single-pixels are independently activated with a focused light pattern; the photovoltage generated largely remains localised within the pixel lateral boundaries, even at high irradiance levels, thus ensuring the absence of electrical crosstalk between the pixels. POLYRETINA delivers a capacitive-like photovoltage optimal for network-mediated activation of the RGCs from the epiretinal side^45^, provided that long (e.g. 10-ms) light pulses are used. The TiN coating also allows single-pixel stimulation at irradiance levels, largely below the MPE limit (8.32 mW mm^−2^ for 10-ms pulses at 565 nm with an illumination rate of 5 Hz).

Conjugated polymers were introduced in retinal stimulation as continuous films directly interfaced with the retina^56,57^ and already proved to be effective in vivo to restore visual acuity in blind rats^14^. In principle, a continuous film might be advantageous compared to discrete electrodes since the fixed arrangement of the electrodes limits the spatial resolution of the stimulation. However, focused stimulation with continuous films is possible only with materials having low carrier mobility and lifetime, such as conjugated polymers^16^. A similar approach is now proposed with other materials like TiO_2_ nanotubes^58^ and Au-TiO_2_ nanowires^18^. However, while the use of a continuous film is interesting for small implants, it becomes more challenging for wide-field implants for the restoration of a large visual angle. Inorganic materials might not be easily fabricated on conformable materials since they often require high-temperature processes. Polymers can be deposited on a large area, but they would immediately crack and eventually delaminate once stretched over a spherical surface like POLYRETINA. Therefore, pixels must be fabricated and protected with rigid elements (i.e. SU-8 platforms) to preserve their mechanical integrity, thus forcing a fixed geometry. Once pixels are introduced for mechanical protection, the coating with metallic cathodes (i.e. Ti/TiN) is convenient because it enhances the stimulation efficiency.

When independent pixels stimulate the highly interconnected and spatially organised network of the retina, an increase in the stimulation resolution may not necessarily translate into an equivalent improvement of the RGC response resolution^41,59,36,17^. First, we assessed the RGC RFs under photovoltaic stimulation: 77% of the stimulated RGCs had an average photovoltaic RF diameter slightly larger than the pixels pitch (153.7 ± 26.1 µm), while 23% of the RGCs exhibit a wider photovoltaic RF (335.5 ± 49.3 µm). The two main clusters of photovoltaic RFs we found can be related to two different sizes of natural RFs belonging to diverse RGC types. The small photovoltaic RFs corresponds to the majority of the natural functional RFs and dendritic trees diameters reported in both wild-type and rd10 mice^60,61,51,49^. However, a more systematic and exhaustive classification of functional RFs revealed that several α-RGCs types exhibit RF diameters varying between 180 and 400 µm^62,63^. The large photovoltaic RFs had an average diameter matching with the dendritic field diameter of large α-RGCs^63^. Second, we demonstrated that adjacent pixels could stimulate a single RGC through different portions of its RF. As a consequence, when consecutively stimulated by two adjacent pixels, the RGC exhibit a marked response to the pixel reversal. The decay of RGC spiking response that is observed during repeated network-mediated stimulation originates from the chemical and kinetical properties of BCs, which desensitise rapidly after electrical stimulation^53,54,64^. The rebound of the RGC activity at the pixel reversal originates from the stimulation of an adjacent pool of not-desensitised BCs. These results suggest that POLYRETINA enables a sub-RF response resolution for RGCs having a dendritic tree larger than the pixel’s pitch. However, the illumination of a single-pixel is expected to induce ML activity in more than one RGC, ideally in all the RGCs whose RFs overlap in the area activated by the illuminated pixel. The indirect stimulation of RGCs, even if non selective for the specific cell type, may preserve the parallel retinal processing, thus providing a more naturalistic activation of the retina.

It is well-founded to assume that the variety of photovoltaic RFs we observed originate from the variety of natural RFs of the mouse model. The peculiarity of mice retina compared to the human one is to have large and rather homogeneous RFs despite eccentricity, with the RGC topography being more specific in a dorsoventral axis, without any region devoted to high visual acuity^65,66^. Most of the natural RFs are equal or larger than the pitch of the photovoltaic pixels within POLYRETINA^62^. As a consequence, the network-mediated photovoltaic RFs and the response resolution are firstly limited by the natural convergence of BCs and only secondly by the pixel density. However, in perspective of clinical applications, it should be noted that the dendritic fields of the RGCs in the human fovea are substantially smaller: 5 to 12 µm for midget cells and 30 to 40 µm for parasol cells^67,68^. The pixel density would thus determine the theoretical resolution limit in the fovea and parafoveal retina. The theoretical acuity limit that can be achieved with 120-µm spaced pixel array is 20/480^69^. This value places POLYRETINA in the upper intermediate level between Argus® II and PRIMA/Alpha-AMS ones, a borderline resolution range for faces and emotions recognition^70^. Nevertheless, such a form of artificial vision may be valuable for a more reliable obstacle recognition and ambulation^20,71^. The primary difference between POLYRETINA and the aforementioned implants is its large visual angle, which has an impact on the perceived field as well as on the peripheral resolution. The combination of both visual acuity and visual angle is recognised as a crucial need to map and interact with one’s environment, having consequences on the layout space understanding, walking distance evaluation, identify-and-reach tasks, spatial cognition and attention^72,73^. The legal definition of blindness in the United States of America and most European countries does not only take into account the foveal acuity (worse than 20/200) but also the visual angle (smaller than 20 degrees), because of its critical role in the naturalistic perception of complex scenes, movements and objects. Indeed, self-orienting task and free mobility in a moving environment require the rapid detection of movements and luminance changes from the entire visual field. Motion-sensitive magnocellular-projecting parasol cells can provide useful depth and motion information from the peripheral retina^74,75,67,76^. Also, fine perception, such as visual hyperacuity, is not only a mechanistic print of the retina activation map but relies on the joint perception of features from the central and the peripheral retina^77,78^.

In humans, contrarily to mice, the size of RFs and the arborisation of both midget and parasol cells increases with eccentricity^67,68,79^. The POLYRETINA prosthesis covers 43 degrees, about 11 to 13% of the retinal surface^80,81^, i.e. the fovea, the parafovea, the perifovea, and up to 6-7 mm away from the fovea in the mid-peripheral retina. The dendritic tree of perifoveal parasol cells vary between 150 and 270 µm and of midget cells between 25 and 70 µm; while they respectively vary between 175 and 310 µm, and 50 and 120 µm in the mid-peripheric retina^67^. Outside of the fovea, both cell types have RF diameters larger or equal to the pixel pitch of POLYRETINA. According to the photovoltaic RFs measured in the mouse model, in such case, the prosthetic resolution is limited by the dendritic arborisations. Altogether, nearly 40% of the human RGCs, mostly parafoveal and mid-peripheral parasol cells but also mid-peripheral midget cells could be stimulated with a resolution matching their physiological RFs.

The restoration of a large visual field with appropriate resolution in the parafoveal mid-peripheral retina represents a leap forward for retinal prostheses. The stimulation of parafoveal to mid-peripheric motion-sensitive cells, as well as perifoveal chromatic and contrast-sensitive midget cells, could facilitate ambulation and motion detection, and be of great help to reduce the need for head scanning and space decomposition, which is a major obstacle today during daily activities.

### Outlook

Our results demonstrated that POLYRETINA could achieve a high spatial resolution in retinal stimulation, together with a wide visual angle, which would be a substantial step forward for artificial vision. However, several steps are still required before considering POLYRETINA for a clinical trial.

One of the present limits of photovoltaic retinal prostheses based on conjugated polymers is the use of semiconducting materials absorbing light in the visible spectrum (e.g. P3HT). The use of visible light for prosthetic activation is not optimal due to the possible activation of remaining photoreceptors in patients with some residual vision. Moreover, the high irradiance levels required to activate POLYRETINA might be perceived even in blind patients without residual vision. Novel conjugated polymers with shifted sensitivity in the far-red and near-infrared could be exploited to overcome this problem^82,83^. Recent results reported the possibility to use a red-shifted polymer in neural interfaces^84^ and retinal prostheses^85^.

Another aspect to be further considered for clinical trials is thermal safety. The MPE was calculated based on retinal damage, but it is not the only element to be considered for safety. According to the thermal safety standard for active implantable medical devices (ISO 14708-1:2014 / EN 45502-1:1997), the maximum temperature rise on the surface of the implant should not exceed 2 °C above the normal surrounding temperature. Thermal simulations for POLYRETINA operating at the MPE already verified such requirement^16^. However, since a light beam is projected into the pupil during photovoltaic stimulation, it might be transiently focused on the iris during involuntary eye movements. In such a case, the temperature of the iris should not increase more than 2 °C to ensure the safe operation of the device. We performed a finite element analysis simulation based on the worst scenario in which the full beam at the MPE is projected onto the iris for a prolonged period. According to the calculated MPE, 47.90 mW at 565 nm could enter the pupil for chronic exposure. Given a constricted pupil of about 3 mm (as considered in the safety standard^86^) and a light beam reduced to a spot of 2 mm in diameter (to avoid beam clipping), the resulting irradiance at the iris plane is 15.25 mW mm^−2^. Due to the axial symmetry of the thermal simulation, we modelled the iris as a continuous tissue without pupil (**Fig. 7a**). The temperature in the iris increased by 13.58 °C after 150 s of continuous illumination at the MPE (**Fig. 7b**, red line), which is largely above the safety limit of 2 °C. In order to keep the thermal increase in the iris below 2 °C, the irradiance should be reduced to 2.25 mW mm^−2^ (**Fig. 7b**, green line). If the irradiance at the iris plane is limited to 2.25 mW mm^−2^, this will correspond to a total of 7.07 mW chronically entering from the pupil. At the retinal level, this irradiance would correspond to a maximum of 1.1 mW mm^−2^ for 10-ms pulses, 5 Hz repetition rate, and an illuminated area of 127.97 mm^2^ (43 degrees), which is still above the irradiance threshold for single-pixel stimulation of RGCs. It is though unlikely that the beam remains statically focused on the same area of the iris for 150 s, as the continuous eye movements would spread the light beam over a larger area and reduce its thermal impact. Therefore, we quantified the time needed to reach 2 °C with a stable beam (**Fig. 7c**). At 15.25 mW mm^−2^, an increase of 2 °C is reached after 460 ms, and the time increases by decreasing the irradiance at the iris plane: it is reasonable to consider that fast saccadic movements will reduce the thermal impact. Moreover, eye-tracking sensors embedded in modern virtual reality glasses provide tracking at 120 Hz, thus allowing real-time adjustment of the beam based on the eye gaze. Under a working hypothesis of 10-ms pulses and 5 Hz of repetition rate, each pulse is delivered every 200 ms, and the eye tracker will have enough time to correct the projection system by using a steering mirror or, in the worst case, to close the beam to preserve the iris. Again, a shift towards near-infrared conjugated polymers will be beneficial for thermal safety at the iris.

**Figure 7.**
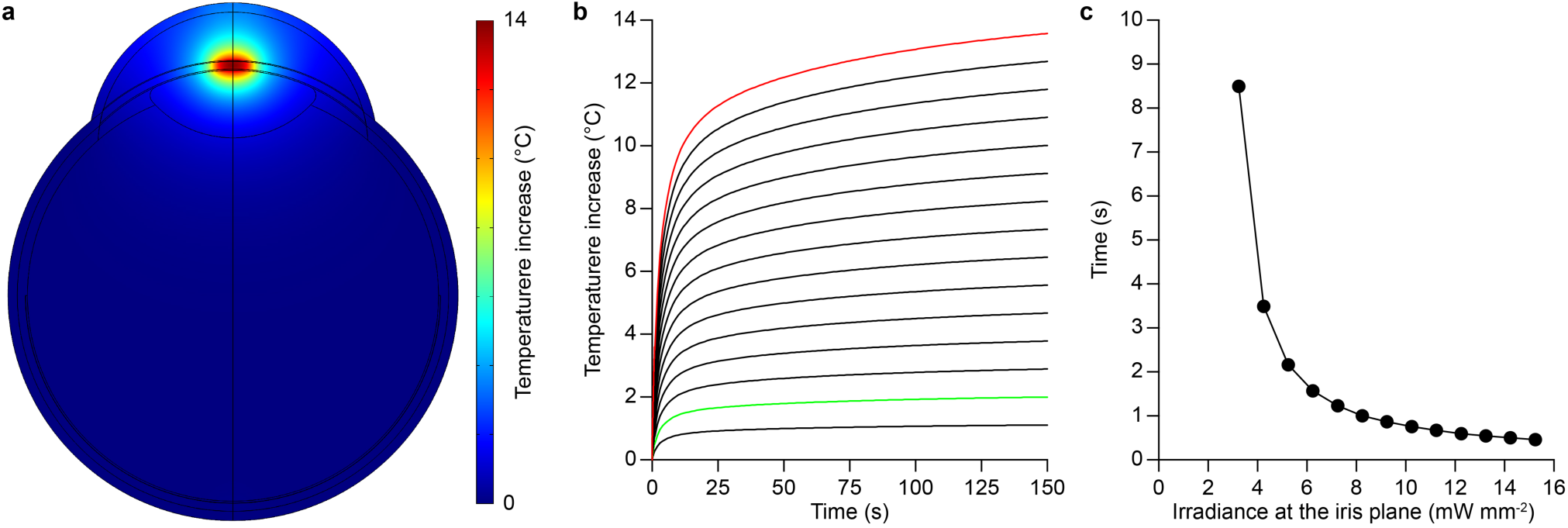
Thermal simulations at the iris plane. **a**, Temperature increase at the iris plane in the modelled eye after 150 s of continuous illumination at the MPE (565 nm, 15.25 mW mm^−2^). **b**, Quantification of the temperature increase at the iris plane during 150 s of continuous illumination for various irradiance levels: 1.25, 2.25, 3.25, 4.25, 5.25, 6.25, 7.25, 8.25, 9.25, 10.25, 11.25, 12.25, 13.25, 14.24 and 15.25 mW mm^−2^. The red line corresponds to 15.25 mW mm^−2^ and the green line to 2.25 mW mm^−2^. **b**, Quantification of the time to reach a thermal increase of 2 °C as a function of the irradiance at the iris plane for continuous illumination at 565 nm.

Last, the POLYRETINA’s safety and efficacy should be validated in preclinical trials in vivo.

## Methods

### Mechanical simulations

Finite element analysis simulations were performed in Abaqus/CAE 6.14, using a three-dimensional deformable shell (photovoltaic interface) moving against a static spherical solid (hemispherical dome) to create a full hard contact. The edges of the shell were clamped to move only in the vertical direction toward the solid dome. The surface roughness and intrinsic thin-film stresses arising from deposition techniques were not considered in the simulation. The shell was constructed using the parameters listed in **Tab. 1**.

**Table 1.**
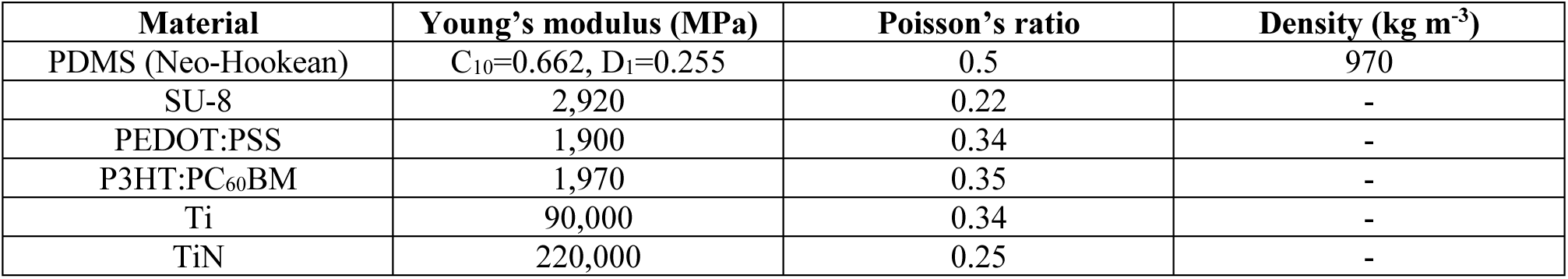
Mechanical simulations. List of parameters used for the construction of the deformable shell. Apart from PDMS, the behaviours of the other materials were considered isotropic elastic. PDMS: polydimethylsiloxane; PEDOT: poly(3,4-ethylenedioxythiophene); PSS: poly(styrenesulfonate); P3HT: regioregular poly(3-hexylthiophene-2,5-diyl); PC_60_BM: [6,6]-phenyl-C61-butyric acid methyl ester; Ti: titanium; TiN: titanium nitride. The values for the Young’s modulus and Poisson’s ratio of the used materials and the hyperelastic coefficients for PDMS were taken from the following references^87–93^.

### Thermal model

COMSOL Multiphysics 5.3 was used with the Bioheat module and the General PDE module for the heat transfer and Beer-Lambert light propagation. A uniform beam with a diameter of 2 mm (565 nm) was used as the illumination source. The eye model was built with several spheres representing each component (cornea, aqueous humour, lens, iris anterior border layer, iris stroma, iris pigmented epithelium, vitreous humour, retina, retinal pigmented epithelium, choroid, and sclera). The iris was simulated as a continuous film completely covering the pupil, while the light beam was projected on the iris, centred to the pupil location. All the parameters used in the model are listed in **Tab. 2**.

**Table 2.** Eye parameters used for the eye model. The parameters were obtained from references^94–100^. The heat capacity, thermal conductivity, density, perfusion rate and self-heat of the iris anterior border layer, stroma, and pigment epithelium were taken respectively from the cornea, choroid and retinal pigment epithelium, due to their biological similarity.

| Material | Thickness | Heat Capacity | Thermal Conductivity | Density | Absorption (565 nm) | Perfusion rate | Self-Heat |
| --- | --- | --- | --- | --- | --- | --- | --- |
|  | μm | J kg<sup>-1</sup> K<sup>-1</sup> | W m<sup>-1</sup> K<sup>-1</sup> | Kg m<sup>-3</sup> | m<sup>-1</sup> | s<sup>-1</sup> | W m<sup>-3</sup> |
| Aqueous humour | 3100 | 3997 | 0.58 | 1000 | 0.025 | 0 | 0 |
| Choroid | 430 | 3840 | 0.53 | 1050 | 15000 | 0.0091 | 10000 |
| Cornea | 500 | 4178 | 0.58 | 1050 | 51 | 0 | 0 |
| Lens | 3600 | 3000 | 0.4 | 1050 | 2.5 | 0 | 0 |
| Retina | 100 | 3680 | 0.565 | 1000 | 400 | 0 | 0 |
| Retinal pigment epithelium | 10 | 4178 | 0.603 | 1050 | 110000 | 0 | 0 |
| Sclera | 500 | 4178 | 0.58 | 1000 | 590 | 0 | 0 |
| Vitreous humour | / | 3997 | 0.6 | 1000 | 0.025 | 0 | 0 |
| Iris anterior border layer | 50 | 4178 | 0.58 | 1050 | 5470 | 0 | 0 |
| Iris stroma | 400 | 3840 | 0.53 | 1050 | 2750 | 0.0091 | 10000 |
| Iris pigment epithelium | 70 | 4178 | 0.603 | 1050 | 100000 | 0 | 0 |

### Chips micro-fabrication

Samples were fabricated on 20 × 24 mm^2^ glass substrates (2947-75×50, Corning Incorporated) cleaned by ultrasonication in acetone, isopropyl alcohol, and deionised water for 15 min each and then dried with a nitrogen gun. PEDOT:PSS (PH1000, Clevios Heraeus) was mixed to 0.1 v/v% (3-glycidyloxypropyl)trimethoxysilane (440167, Sigma Aldrich), filtered (1 μm PTFE filters), and then spin-coated at 3000 rpm for 40 seconds on each chip. Subsequent annealing at 115 °C for 30 minutes was performed. The preparation of the bulk heterojunction was performed in a glove box under nitrogen atmosphere. 20 mg of P3HT (M1011, Ossila) and 20 mg of PC_60_BM (M111, Ossila) were dissolved in 1 mL of anhydrous chlorobenzene each and let stirring overnight (16 hr) at 70 °C. The solutions were then filtered (0.45 μm PTFE filters) and blended (1:1 v:v). The P3HT:PC_60_BM blend was spin-coated at 1000 rpm for 45 seconds. Subsequent annealing at 115 °C for 30 min was performed. Titanium and titanium nitride cathodes were deposited by direct-current (Ti) and radio frequency (TiN) magnetron sputtering using a shadow mask. The polymer patterning step was obtained by exposing the chips to oxygen plasma. A plastic reservoir was then attached to the sample using PDMS as an adhesive.

### Measure of photo-voltage and photo-current

Samples were placed on a holder, and each electrode was sequentially contacted. A platinum wire immersed in physiological saline solution (NaCl 0.9 %) was used as a counter electrode. 10-ms light pulses were delivered by a 565-nm LED (M565L3, Thorlabs) focused at the sample level. Photo-voltage and photo-current were measured using respectively a voltage amplifier (1201, band DC-3000 Hz, DL-Instruments) and a current amplifier (1212, DL-Instruments). Data sampling (40 kHz) and instrument synchronisation were obtained via a DAQ board (PCIe-6321, National Instruments) and custom-made software. Data analysis was performed in MATLAB (MathWorks). When evaluating the photo-current density generated by the interface, the area of the connecting line exposed to light was also considered.

### POLYRETINA micro-fabrication

Photovoltaic interfaces were fabricated on silicon wafers. A thin sacrificial layer of poly(4-styrenesulfonic acid) solution (561223, Sigma-Aldrich) was spin-coated on the wafers (1500 rpm, 40 s) and baked (120 °C, 15 min). Degassed PDMS pre-polymer (10:1 ratio base-to-curing agent, Sylgard 184, Dow-Corning) was then spin-coated (1000 rpm, 60 s) and cured in the oven (80 °C, 2 hr). After surface treatment with oxygen plasma (30 W, 30 s), a 6-µm thick SU-8 (GM1060, Gersteltec) layer was spin-coated (3800 rpm, 45 s), soft-baked (130 °C, 300 s), exposed (140 mJ cm^−2^, 365 nm), post-baked (90 °C, 1800 s; 60 °C, 2700 s), developed in propylene glycol monomethyl ether acetate (48443, Sigma-Aldrich) for 2 min, rinsed in isopropyl alcohol and dried with nitrogen. After surface treatment with oxygen plasma (30 W, 30 s), a second layer of degassed PDMS pre-polymer (10:1) was spin-coated (3700 rpm, 60 s) and cured in the oven (80 °C, 2 hr). PEDOT:PSS and P3HT:PC_60_BM were prepared and deposited as described before. Titanium and titanium nitride cathodes were deposited by direct-current (Ti) and radio frequency (TiN) magnetron sputtering using a shadow mask aligned with the SU-8 pattern. After the patterning of polymers by oxygen plasma, the encapsulation layer of degassed PDMS pre-polymer (5:1 ratio) was spin-coated (4000 rpm, 60 s) and cured in the oven (80 °C, 2 hr). Photolithography and PDMS dry etching were performed to expose the cathodes. The wafers were then placed in deionised water to allow for the dissolution of the sacrificial layer and the release of the photovoltaic interfaces. The floating membranes were finally collected and dried in air. The hemispherical PDMS domes were fabricated using a milled PMMA mould, filled with PDMS pre-polymer (10:1), which was then degassed and cured in the oven (80 °C, 2 hr). The supports were released from the moulding parts and perforated with a hole-puncher (330-µm in diameter) at the locations dedicated to the insertion of retinal tacks. The released photovoltaic interfaces were clamped between two O-rings and, together with the hemispherical domes, were exposed to oxygen plasma (30 W, 30 s). The activated PDMS surfaces were put in contact and allowed to uniformly bond thanks to radial stretching of the fixed membrane. The excessive PDMS used to clamp the array was removed by laser cutting.

### Atomic force microscopy

AFM images and roughness measurements were obtained with a Bruker Dimension icon microscope and scanasyst-air Si tips. Images (500 nm x 500 nm) were plotted and the surface area was calculated with NanoScope analysis 1.9 software.

### Kelvin Probe Force Microscopy

KPFM characterisation was performed in ambient air conditions with a dimension icon atomic force microscope (Bruker Corporation) using n-doped silicon tips (SCM-PIT-V2, Bruker Corporation) in surface potential, amplitude-modulated imaging mode. KPFM images were collected by repetitively scanning a single 100-nm line under dark and light conditions to measure the surface potential variation. The green LED of a Spectra X illumination system (Emission filter 560/32, Lumencor) was used to illuminate the pixel using an optical fiber and focused onto the pixel (Photo-Conductive accessory, Bruker Corporation). The samples were grounded using a silver paste; however, individual pixels could not be connected to the paste and were therefore floating. The voltage bias was sent to the AFM tip. KPFM images were analysed using Gwyddion 2.36 software. For each image, the average surface potential variation value was obtained by subtracting the surface potential in the dark to the one under illumination (voltage in light – voltage in dark).

### Spatial selectivity measures

Measures of the voltage spread were performed in Ames’ medium (A1420, Sigma-Aldrich) at 32 °C with a glass micropipette (tip diameter about 10 μm) located approximately 2-5 µm from the implant surface. Data were amplified (Model 3000, A-M System), filtered (DC - 1,000 Hz), and digitalised at 30 kHz (Micro1401-3, CED Ltd.). Illumination was carried out on a Nikon Ti-E inverted microscope (Nikon Instruments) using a Spectra X illumination system (Emission filter 560/32, Lumencor). The microscope was equipped with a dichroic filter (FF875-Di01-25×36, Semrock) and a 10x (CFI Plan Apochromat Lambda) objective. The patterning of the light stimulus was carried out using a light patterning system (Polygon 400, Mightex). The light pattern sequences were adjusted in real-time to align the light patterns to the prosthesis pixels (PolyScan, Mightex). After alignment of the illumination pattern onto the POLYRETINA pixels, 10 pulses of 10 ms were delivered at 1 Hz with an irradiance of 22.65 mW mm^−2^. Data analysis was conducted in MATLAB. Voltage peaks above noise level were detected, and their amplitude normalised respect to the central pixel value.

### Electrophysiology

Animal experiments were conducted according to the animal authorizations GE3717 issued by the Département de l’Emploi, des Affaires sociales et de la Santé (DEAS), Direction Générale de la Santé of the République et Canton de Genève (Switzerland). Both male and female rd10 mice were used (**Tab. 3**). Mice were kept in a 12 h day/night cycle with access to food and water ad libitum. White light (300 ± 50 lux) was present from 7 AM to 7 PM and red light (650-720 nm, 80-100 lux) from 7 PM to 7 AM.

Retinas from inbred Rd10 mice colony were explanted in normal light conditions after the animals were sacrificed by injection of Sodium Pentobarbital (150 mg kg^−1^). After eye enucleation, retinas were dissected in carboxygenated (95 % O_2_ and 5 % CO_2_) Ames’ medium (A1420, Sigma-Aldrich) and transferred to the microscope stage for stimulation and recording. Retinas were placed with the retinal ganglion cells facing down on the prosthesis. Recordings were performed in dim light at 32 °C with a sharp metal electrode (PTM23BO5KT, World Precision Instruments), amplified (Model 3000, A-M System), filtered (300-3000 Hz), and digitalized at 30 kHz (Micro1401-3, CED Ltd.). Illumination was carried out on a Nikon Ti-E inverted microscope (Nikon Instruments) using a Spectra X illumination system (Emission filter 560/32, Lumencor). The microscope was equipped with a dichroic filter (FF875-Di01-25×36, Semrock) and a 10x (CFI Plan Apochromat Lambda) objective. The patterning of the light stimulus was carried out using a light patterning system (Polygon 400, Mightex). The light pattern sequences were real-time adjusted to align the light patterns to the prosthesis pixels (PolyScan, Mightex). After alignment of the illumination pattern onto the POLYRETINA pixels, for each retinal ganglion cell, 10 pulses of 10 ms were delivered at 1 Hz for each illumination condition. Spike detection and sorting were performed by threshold detection using the Matlab-based algorithm Wave_clus^101^ and further data processed in MATLAB. An exclusion period of ± 1 ms around light onset and offset was applied to avoid artefact misclassification.

### Optical safety

Retinal damage upon light exposure can occur because of three main factors: photo-thermal damage, photo-chemical damage, and thermo-acoustic damage^86^. In ophthalmic devices, Maxwellian illumination is used where the incident illumination occupies a fraction of the pupil (no overfilling). For continuous illumination, the MPE could be controlled by the photo-thermal (MPE_T_) or photo-chemical damage (MPE_C_), calculated in W according to equations (1) and (2), respectively.

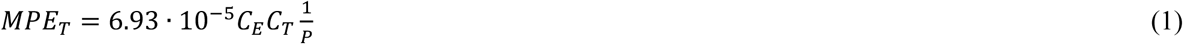

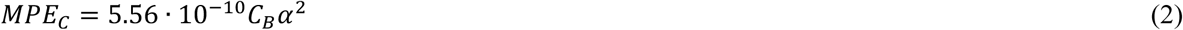

For POLYRETINA, the visual angle *α* is calculated according to equation (3), and the exposed area according to equation (4), in which *d* = 13.4 mm is the diameter covered by the active area and *f* = 17 mm is the eye’s focal length.

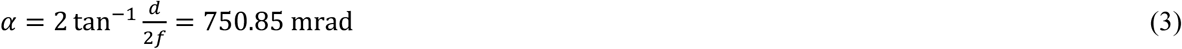

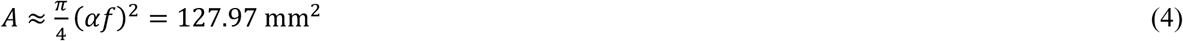

For λ = 565 nm, both limits apply and *C*_*E*_ = 6.67 · 10^−3^*α*^2^; *C*_*T*_ = 1; *P* = 5.44; *C*_*B*_ = 10^0.02.(*λ* −450)^. The limits are MPE_T_ = 47.90 mW and MPE_C_ = 62.54 mW. Therefore, the limiting factor is MPE_T_ which results in 47.90 mW entering the pupil, and corresponds to 374.3 µW mm^−2^ for an exposed area of 127.97 mm^2^. However, POLYRETINA operates with pulsed illumination. With pulses of 10 ms and duty cycle of 20, 10, or 5 % (respectively for 20, 10, or 5 Hz), the MPE is increased to 1.87, 3.74, or 7.48 mW mm^−2^ respectively^85^. In addition, a previous thermal model showed that at 565 nm and over the broad range of irradiance levelss the temperature increase in the retina is reduced by 11 % with POLYRETINA^16^. Therefore, the MPE could be increased to 2.08, 4.16, or 8.32 mW mm^−2^ respectively for 20, 10, or 5 Hz.

### Statistical analysis and graphical representation

Statistical analysis and graphical representation were performed with Prism (GraphPad Software Inc.) and MATLAB. The normality test (D’Agostino & Pearson omnibus normality test) was performed in each dataset to justify the use of a parametric or non-parametric test. In each figure p-values were represented as: * p < 0.05, ** p < 0.01, *** p < 0.001, and **** p < 0.0001.

## Data availability

The authors declare that all other relevant data supporting the findings of the study are available in this article and in its supplementary materials. Access to our raw data can be obtained from the corresponding author upon reasonable request.

## Acknowledgement

We would like to acknowledge the Center of Micronanotechnology at École polytechnique fédérale de Lausanne and The Neural Microsystems Platform at the Wyss Center for Bio and Neuroengineering for their support. We would like to acknowledge Jacob Thorn (at École polytechnique fédérale de Lausanne) for English editing. This work was supported by École polytechnique fédérale de Lausanne, Medtronic, Fondation Pierre Mercier pour la Science, Velux Stiftung (Project 1102), Fondation Pro Visu and Gebert Rüf Stiftung (Project GRS-035/17).

## Author contributions

N.A.L.C. designed, performed and analysed spatial selectivity measurements and electrophysiological experiments, and wrote the manuscript. M.J.I.A.L. designed, fabricated, and characterised the prosthesis, and performed mechanical and thermal simulations. D.G. designed and led the study, and wrote the manuscript. All the authors read and accepted the manuscript.

